# Tyloxapol inhibits ESX-1 secretion in *Mycobacterium marinum*

**DOI:** 10.64898/2026.02.02.703260

**Authors:** Owen A. Collars, Patricia A. Champion

## Abstract

Mycobacteria have a hydrophobic cell envelope that makes uniform growth in liquid culture challenging. Non-ionic detergents including Tween-80 and tyloxapol are commonly added to media when culturing mycobacterial species in the laboratory. Tyloxapol was reported to exhibit anti-tuberculous activity during animal infection with *M. tuberculosis* in the 1950s. In the 1980s, microscopy studies suggested that tyloxapol impacted the interaction between *M. tuberculosis* and the phagosomal membrane, preventing mycobacterial access to the cytoplasm. It is now known that the ESX-1 Type VII secretion system mediates the interaction between pathogenic mycobacteria and the phagosomal membrane. *Mycobacterium marinum* is a pathogenic mycobacterial species that has been widely used to understand the molecular mechanisms and host responses to the ESX-1 system. The hemolytic activity of *M. marinum* allows the study of ESX-1 lytic activity outside of the context of a host cell. We found that tyloxapol inhibits the hemolytic activity of *M. marinum* in a concentration dependent manner. The addition of 100-fold less tyloxapol than commonly used for mycobacterial growth differentially inhibits the production and secretion of ESX-1 substrates required for lytic activity. Our findings directly impact how the field interprets data from studies where *M. marinum*, and potentially other mycobacterial species were grown in tyloxapol. Our findings may explain the original observations linking tyloxapol to anti-tuberculosis activity.

**Author Summary:** Tuberculosis, which is caused by *Mycobacterium tuberculosis*, is one of the world’s deadliest diseases. We lack a clear understanding of how *M. tuberculosis* and related mycobacterial species cause disease. In the 1950’s, it was reported that treating *M. tuberculosis* infected animals with tyloxapol improved the survival and in some cases protected the animals from death. Tyloxapol is a detergent that is commonly added to mycobacterial cultures to promote dispersed growth in the laboratory. Later studies suggested that tyloxapol altered the interaction between *M. tuberculosis* and the phagosomal membrane during macrophage infection. The ability to escape the phagosome is essential for mycobacteria to cause disease, and is mediated by a Type VII protein secretion system, ESX-1. Using *M. marinum*, a well-established model for understanding the molecular mechanisms of ESX-1 secretion, we show that tyloxapol used at more than 100-fold less than what is commonly used to grow mycobacteria in the lab, inhibits ESX-1 secretion. Our findings have widespread implications on how we interpret our findings as a field, and may explain why tyloxapol impacted *M. tuberculosis* infection of both animals and macrophages. Our study also indicates that tyloxapol can be used as a tool to understand the molecular mechanisms of ESX-1 protein secretion.

## Introduction

*Mycobacterium marinum*, which causes a tuberculosis-like infection of ectothermic animals (1), is a widely accepted model for understanding mycobacterial pathogenesis, and in particular for studying protein secretion (2, 3). Type VII secretion systems promote mycobacterial and Gram-positive pathogenesis and physiology (4). ESX-1 is a conserved Type VII system in *M. marinum* and in *M. tuberculosis* (5-8). Early during macrophage infection, mycobacterial pathogens are phagosomal (9). The ESX-1 system disrupts the phagosomal membrane allowing bacterial translocation to the cytoplasm, which is required for disease progression (10-12).

The non-ionic detergents Tween-80 and tyloxapol are used in culturing mycobacterial cells to prevent aggregation during *in vitro* growth. In the 1950s, tyloxapol was reported to exhibit anti-tuberculous activity during animal infection with *M. tuberculosis* (13-16). Microscopy-based studies in the 1980s suggested that tyloxapol impacted the interaction between *M. tuberculosis* and the phagosomal membrane, preventing mycobacterial access to the cytoplasm (17, 18). Based on these studies, we hypothesized that tyloxapol directly inhibited ESX-1 activity.

## Results

### Tyloxapol inhibits hemolytic activity in *M. marinum* in a concentration dependent manner

*M. marinum* lyses red blood cells in an ESX-1 and contact-dependent manner (8), allowing the study of ESX-1 lytic activity in the absence of a host cell. To test if growth in tyloxapol impacted hemolytic activity, we grew the WT and Δ*eccCb*_*1*_ *M. marinum* strains in 7H9 media with 0.1% Tween-80 (Fig. 1A). EccCb_1_ is required for ESX-1 secretion (5, 19, 20). We washed the cells and cultured the strains in 7H9 media with either 0.1% Tween-80 or 0.2% tyloxapol at pH 6.8 for 24 hours. The cells were washed to remove the detergents and the buffered media, and then resuspended in PBS to perform the hemolysis assays. As shown in Fig. 1B, the WT strain was significantly more hemolytic than the Δ*eccCb*_*1*_ strain following growth with Tween-80 at pH 6.8. However, there was no significant difference in the hemolytic activity of the WT and the Δ*eccCb*_*1*_ strains following growth with tyloxapol at pH 6.8. From these data, we conclude that growth in tyloxapol inhibited the ESX-1-dependent hemolytic activity of *M. marinum*.

**Figure 1.**
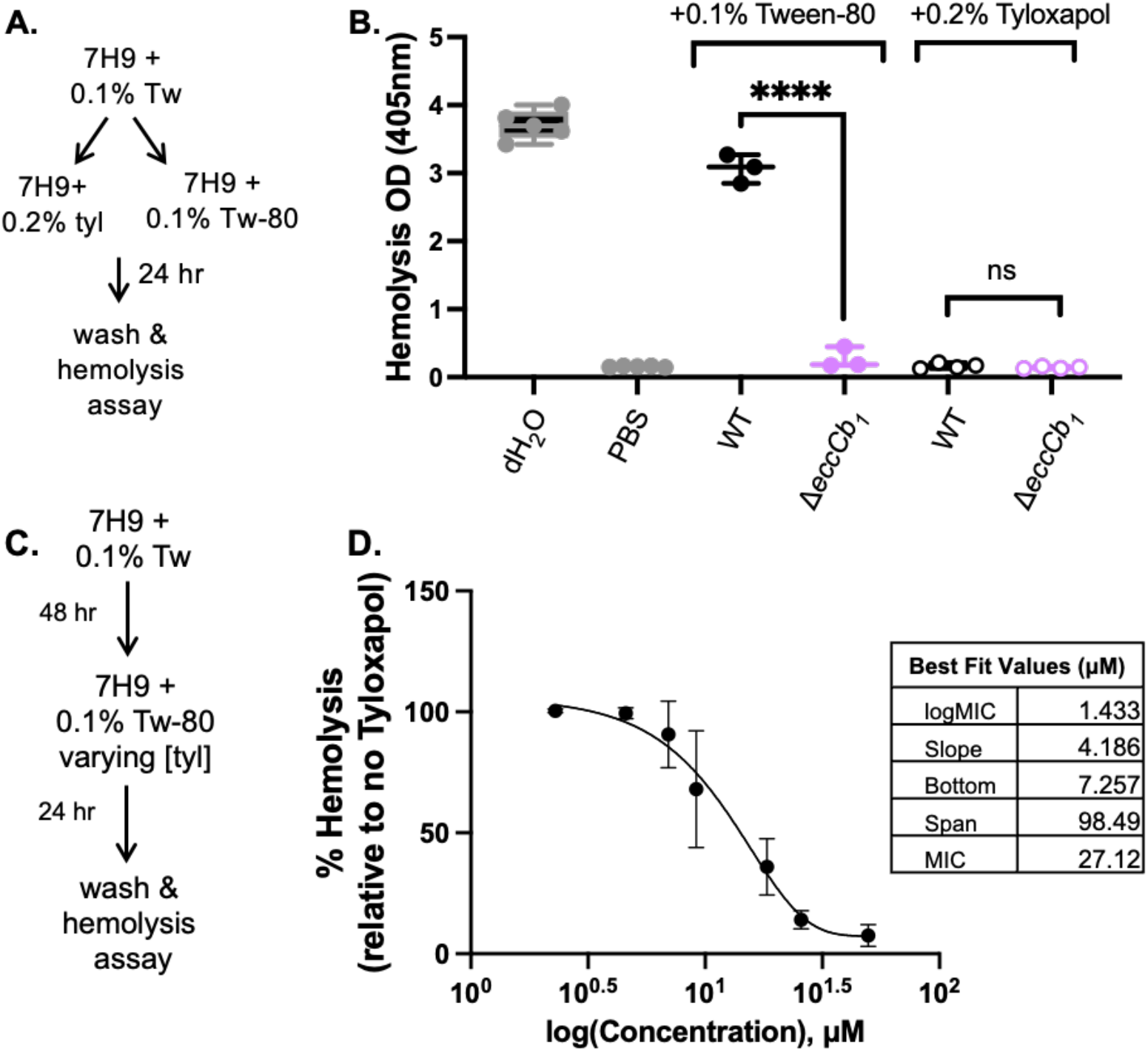
Tyloxapol inhibits ESX-1-dependent hemolytic activity in a concentration dependent manner. (**A)** Schematic of growth conditions for panel B. **(B)** Hemolytic activity of *M. marinum* grown in 0.1% Tween-80 or 0.2% tyloxapol for 24 hours. Each data point represents a biological replicate consisting of three technical replicates. Significance was determined using a one-way ordinary ANOVA (*P*<0.0001) followed by a Tukey’s multiple comparison test **** *P*<0.0001. **(C)** Schematic of growth conditions for panel D. **(D)** Tyloxapol inhibition curves of hemolytic activity. Bacteria were grown in the presence of 0.1% Tween-80 with varying amounts of tyloxapol from 0.000625% through 0.01355%. Individual data points represent the average of three independent experiments. Error bars are the standard deviation. MIC was calculated using a Gompertz equation for MIC determination. Tyloxapol concentration was determined using an estimated molecular weight of 3000 g/mol, which is the median distance between the upper and lower bounds of the polymer lengths reported by Sigma and Santa Cruz Chemicals.

We next tested if the inhibition of hemolysis by tyloxapol was concentration dependent. We grew *M. marinum* in 7H9 buffered to pH 6.8 in 0.1% Tween-80 with varying concentrations of tyloxapol for 24 hours, washed the cells and performed hemolysis assays (Fig. 1C). As shown in Fig. 1D, the MIC of tyloxapol sufficient to block *M. marinum* hemolytic activity was 27.12 μM. The inhibition of *M. marinum* hemolytic activity was concentration dependent.

### Tween-80 catabolism is not a signal for hemolytic activity

Tween-80 is catabolized by mycobacteria into oleic acid, which can be used as carbon source (21-23). Tyloxapol is not catabolized by mycobacteria (24). To rule out that the oleic acid catabolite promoted hemolysis, we supplemented cultures with 200μM oleic acid and measured hemolysis. The addition of oleic acid did not significantly impact the hemolytic activity of the WT strain under the conditions tested (Fig. 2A). We likewise found that the addition of oleic acid did not impact *M. marinum* growth (Fig. 2B). From these data, we conclude that the oleic acid product due to Tween-80 catabolism does not promote *M. marinum* hemolysis.

**Figure 2.**
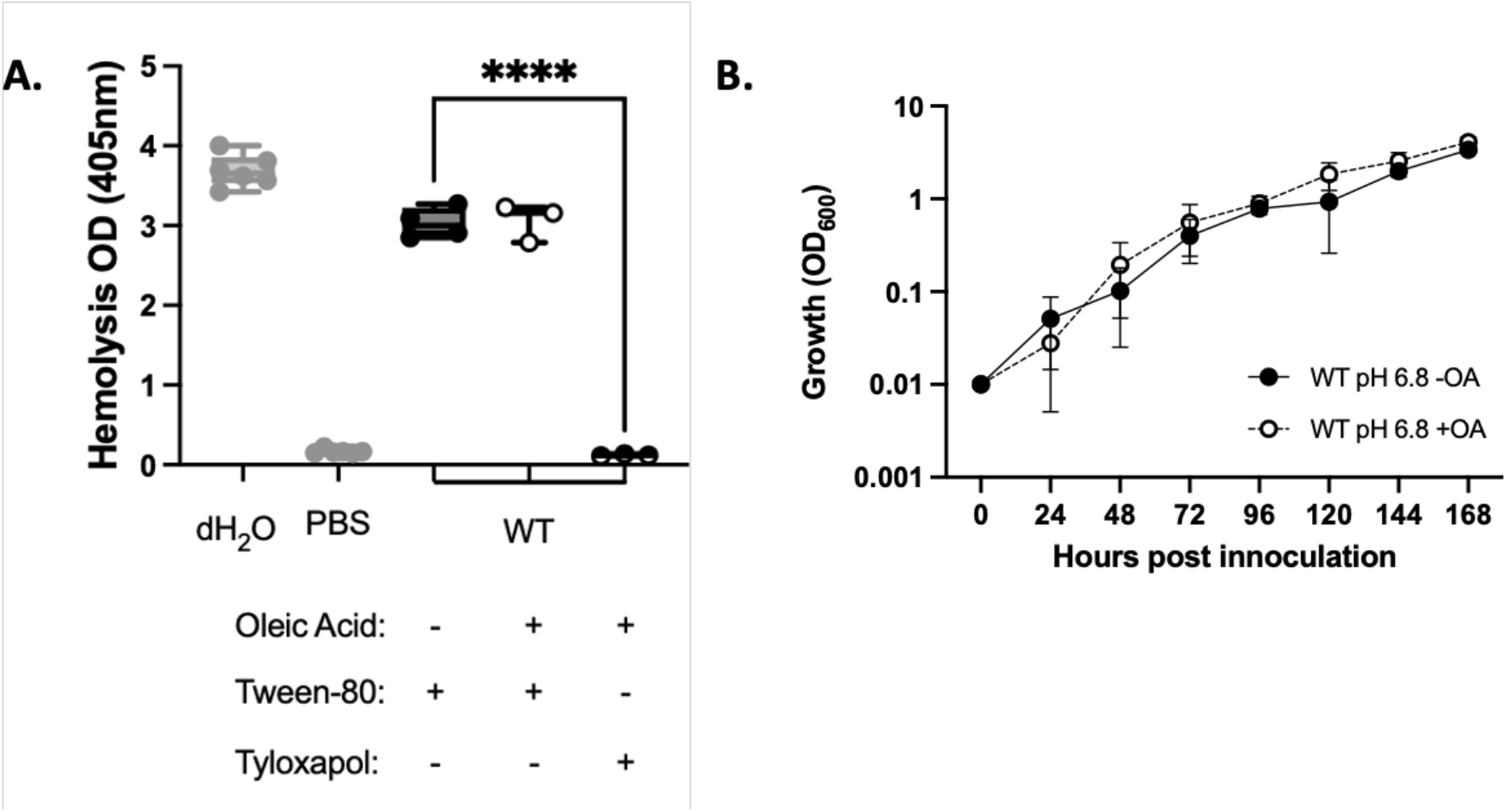
Oleic Acid is not a hemolysis stimulating signal. **(A)** Hemolytic activity of *M. marinum* grown in with 0.1% Tween-80 or 0.2% tyloxapol, with or without 200mM oleic acid for 24 hours. Each data point represents three biological replicates, each including three technical replicates. Significance was determined using a one-way ordinary ANOVA (*P*<0.0001) followed by a Tukey’s multiple comparison test comparing each sample to the WT strain grown in 0.1% Tween-80. **** *P*<0.0001. **(B)** WT *M. marinum* strains in 7H9 media with 0.2% tyloxapol. Strains were grown for 7 days, with growth readings taken every 24 hours by OD_600_. Statistical analysis using a 2way ANOVA followed by a Tukey’s multiple comparison test. The two curves are not significantly different. The y-axis is log_10_ scale.

### Tyloxapol differentially impacts ESX-1 substrate production and secretion

ESX-1 substrates are secreted in a hierarchy under standard laboratory conditions (20). Three groups of substrates undergo ordered secretion by the ESX-1 membrane complex (Fig 3A, light purple). Group I substrates, including the PPE68/MMAR_2894 and EsxA /EsxB (ESAT-6/CFP-10) substrate pairs (red, Fig 3A), are required for Group II substrate secretion (EspB/EspK pair and EspJ, teal) and Group III substrate secretion (EspE/EspF pair, purple). The Group II substrates are required for Group III substrate secretion. The Group III substrates are not required for ESX-1 substrate secretion, but are essential for hemolytic activity and virulence in a macrophage model of infection (20, 25).

**Figure 3.**
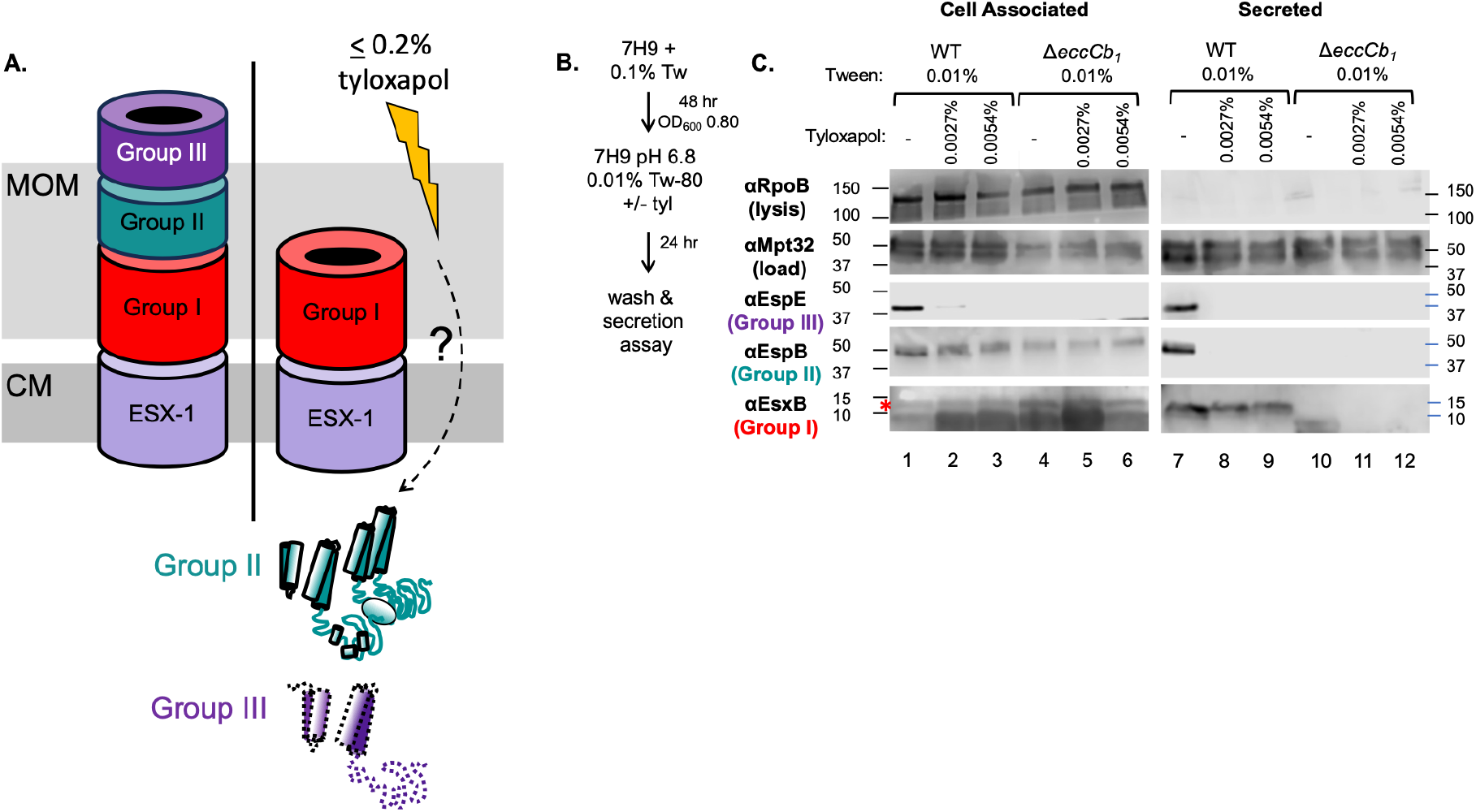
Tyloxapol differentially affects the production and secretion of ESX-1 substrates in *M. marinum*. **(A)** Schematic of ESX-1 secretion assembly. CM= cytoplasmic membrane. MOM= mycolate outer membrane. ESX-1: ESX-1 conserved components including EccA1, EccB_1_, EccCa_1_, EccCb_1_, EccD_1_, EccE_1_, MycP_1_. Group I, II and III substrates are described in the text, as designated in Cronin et al. We propose that tyloxapol is sensed by the mycobacterial cell by an unknown mechanism. Our data supports that Group II substrates are made, but not secreted, while the Group III substrates are not made or secreted **(B)** Schematic of growth conditions for panel C. **(C)** Secretion assay in the presence of 0.0027% and 0.0054% tyloxapol. 30 μg of protein was loaded in each lane. immunoblots are representative of at least three biological replicates. MPT-32 is a protein secreted by the Sec secretion system, and is a loading control for both cell-associated and secreted protein fractions. RpoB is a component of RNA polymerase. RpoB is a loading control for the cell associated fraction, and a lysis control for the secreted protein fraction. The EsxB band is marked by an asterisk in lanes 1-6.

To test how tyloxapol impacted the production and secretion of the ESX-1 substrates, we wanted to mimic the media conditions from the hemolysis assay in Fig. 1. Secreted proteins have historically been isolated from *Mycobacterium* grown in Sauton’s medium (5, 8, 26, 27). We tested if ESX-1 secretion could be detected in the presence and absence of tyloxapol following growth in 7H9. As shown in Fig. 3B, EsxB (Group I), EspB (Group II) and EspE (Group III) were made (lane 1) and secreted from the WT strain (lane 7) grown in 7H9 media supplemented with 0.01% Tween-80. Addition of 0.0027% or 0.0054% tyloxapol to 7H9 supplemented with 0.01% Tween-80 differentially affected the production and secretion for the ESX-1 substrates. Notably, EsxB, was produced (lanes 2 and 3) and secreted from the WT strain (lanes 8 and 9) in the presence of tyloxapol. Although EspB was produced (lanes 2 and 3), EspB was not secreted from the WT strain grown in tyloxapol (lanes 8 and 9). Tyloxapol reduced the level of EspE produced in the WT strain (lanes 2 and 3), and resulted in loss of detectable EspE secretion (lanes 8 and 9). EspE was not detected in the cell associated fraction of the Δ*eccCb*_*1*_ strain due to transcriptional feedback control (28-30). EsxB, EspB and EspE were not secreted from the Δ*eccCb*_*1*_ strain. From these data we conclude that tyloxapol differentially affected the production of the EspB (Group II) and EspE (Group III) substrates, and results in a loss of EspB and EspE secretion, likely explaining the loss of hemolysis of *M. marinum* following growth in tyloxapol.

## Discussion

Detergents are widely used in culturing mycobacterial species in the laboratory. At the time of this manuscript, 223 publications were found in PubMed Central when “*Mycobacterium marinum* and tyloxapol” were searched together (Table 1), with 31 in 2025 alone. The studies that use tyloxapol for growth focus on a range of topics, from fundamental physiology to pathogenesis. Tyloxapol is also exclusively used in media preparations aimed at maintaining virulence lipids production during laboratory growth of *Mycobacterium* (31).

**Table 1.** Number of publications in PubMed Central in response to searches including the species name “and” tyloxapol or Tween-80.

| **Species** | **tyloxapol** | **Tween-80** |
| --- | --- | --- |
| *M. marinum* | 223 | 1342 |
| *M. tuberculosis* | 1154 | 9500 |
| *M. smegmatis* | 747 | 4204 |
| *M. abscessus* | 206 | 1161 |

In this study, we showed that tyloxapol, at levels 100-200x lower than are commonly used in mycobacterial culturing, differentially blocks the production and secretion of ESX-1 substrates from *M. marinum*, abrogating hemolytic activity. It is well established that the Group II and Group III substrates, which are impacted by tyloxapol, are essential for mycobacterial virulence (8, 20, 25, 32-34). We and others have shown that deletion of the ESX-1 system in *M. marinum* or in *M. tuberculosis* alters transcription (28, 35). It is unclear if tyloxapol inhibits the ESX-1 system in other mycobacterial species or the activity of the paralogous ESX systems. Tyloxapol may have additional impacts on mycobacterial physiology, which may alter how we interpret our work as a field.

Our work may explain earlier literature linking tyloxapol to anti-tubercular activity. Beginning in the 1950’s, several independent groups published data supporting that tyloxapol (also named Macrolon, Triton WR-1339) is protective against *M. tuberculosis* infection (13-15, 36). Although tyloxapol is not bactericidal, adding tyloxapol to the growth media of primary macrophages inhibited *M. tuberculosis* growth during infection (37). Likewise, macrophages isolated from tyloxapol-treated mice were more resistant to *M. tuberculosis* infection. Treating mice with tyloxapol prior to or following *M. tuberculosis* infection was protective against death. In fact, the CFUs from lungs and spleens, and time to death from mice infected with *M. tuberculosis* in the presence of tyloxapol were similar to infections with the attenuated BCG vaccine strain (38, 39), which has a natural deletion of several ESX-1 genes (7, 40). The addition of tyloxapol before or after infection of guinea pigs and mice reduced infection or even healed infected animals(16). Microscopy data from the 1980’s suggested that tyloxapol altered the interaction between *M. tuberculosis* and the phagosome membrane (17).

Historically, mycobacterial short term culture filtrates (STCFs) have been generated from cultures grown in Sauton’s medium (26). In *M. smegmatis*, ESX-1 substrates were detected in STCFs from strains grown in Sauton’s medium, but not from 7H9 medium (27). Our study indicates that ESX-1 proteins from *M. marinum* can be detected from STCFs generated from cultures grown in 7H9 medium.

We demonstrate that tyloxapol blocks ESX-1 activity *in vitro* by impacting the secretion of the Group II and Group III substrates, but not Group I substrates, from *M. marinum*. Tyloxapol could affect targeting or translocation of the Group II substrates, effectively stopping ESX-1 from switching from the Group I substrates to secreting the Group II substrates and beyond. The reduced levels of EspE could indicate a feedback mechanism in response to a loss of Group II secretion. Tyloxapol may be a new tool to study the mechanics of ESX-1 secretion. Current studies are focused on identifying the pathways that sense tyloxapol and the response that disrupts ESX-1 secretion.

## Materials and Methods

### Growth and Generation of Bacterial Strains

The *M. marinum* M strain (ATCC BAA-535) and the Δ*eccCb*_*1*_ strain (19) were maintained in Middlebrook 7H9 broth (Sigma-Aldrich, St Louis MO) with 0.5% glycerol and 0.1% Tween-80 (Fisher Scientific, Pittsburgh PA) or on Middlebrook 7H11 agar (Sigma-Aldrich) with 0.5% glycerol and 0.5% glucose at 30°C. To buffer the pH, 7H9 broth was supplemented with 100mM 3-(*N*-morpholino)propanesulfonic acid (MOPS) (ThermoFisher, Waltham, MA) and buffered to pH 6.8 (7H9 pH6.8). Tyloxapol (Chem-Impex, Wood Dale IL) was added as indicated.

### Hemolysis Assay

*M. marinum* were grown in 7H9 broth with 0.1% Tween-80 to mid-log phase. 24 hours prior to the hemolysis assay, the bacteria were sub-cultured into 7H9 pH 6.8 overnight. Where applicable, varying concentrations of tyloxapol as indicated in the figures and text, or 200μM oleic acid (Thermo-Fisher, Waltham MA) was added to cultures. Hemolysis assays were performed exactly as in (20).

### Secretion Assay

*M. marinum* strains were grown in 7H9 media supplemented with glu-cose, glycerol and 0.1% Tween-80 until turbid for ∼5 days. The cultures were diluted to an OD_600_ of 0.8 in 50 mLs of 7H9 pH 6.8 with 0.01% Tween-80, in the presence and absence of tyloxapol as indicated. The cells were grown for 48 hours at 30°C. The culture supernatants and cell associated fractions were prepared as described in (20) except following protein quantification proteins were precipitated in four times volume of acetone (Thermo-Fisher) for one hour. Samples were centrifuged at 14K rpm for 10 minutes at 4 °C. Acetone was removed, protein samples were air dried and resuspended in 16μL of PBS prior to SDS-PAGE. 20µg of protein resolved using 4-20% TGX Gradient Gels (Bio-Rad).

### Immunoblotting

Immunoblots were performed as (41), except they were imaged using a Licor C-Digit digital developer (LICORbio, Lincoln NE) and analyzed using Image Studio 6.1 (LICORbio). The following reagents were obtained through BEI Resources, NIAID, NIH: polyclonal anti-*Mycobacterium tuberculosis* Mpt32 (*Rv1860*, antiserum, rabbit) NR-13807 (1:20,000), polyclonal anti-*Mycobacterium tuberculosis* CFP10 (*Rv3874*, antiserum, rabbit) NR-13801 (1:5,000). The following antibodies were generated by Genscript (Piscataway, NJ): EspE: 1:5,000 dilution, rabbit, epitope: CGQQATLVSDKKEDD based on (34); EspB: 1:1,000 dilution, rabbit, epitope: TKADLEPVNPPKPP based on (32).

## Acknowledgments

This publication was supported by the Institute of Allergy and Infectious Disease of the National Institutes of Health under award numbers R01AI188782 to PAC. The content is solely the responsibility of the authors and does not necessarily represent the official views of the National Institutes of Health. We thank Dr. Matthew Champion for helpful discussion regarding the molecular weight of the tyloxapol polymer.

